# Consistent and scalable monitoring of birds and habitats along a coffee production intensity gradient

**DOI:** 10.1101/2024.07.12.603271

**Authors:** Marius Somveille, Joe Grainger-Hull, Nicole Ferguson, Sarab S. Sethi, Fernando González-García, Valentine Chassagnon, Cansu Oktem, Mathias Disney, Gustavo López Bautista, John Vandermeer, Ivette Perfecto

## Abstract

Land use change associated with agricultural intensification is a leading driver of biodiversity loss in the tropics. To evaluate the habitat-biodiversity relationship in production systems of tropical agricultural commodities, which is critical for certifying and examining the success of biodiversity-friendly agricultural practices, birds are commonly used as indicators. However, consistently and reliably monitoring how bird communities are affected by land use change throughout the annual cycle in a way that can be scalable is challenging using traditional survey methods. In this study, we examined whether the automated analysis of audio data collected by passive acoustic monitoring, together with the analysis of remote sensing data, can be used to efficiently monitor avian biodiversity along the gradient of habitat degradation associated with the intensification of coffee production. Coffee is an important crop produced in tropical forested regions, whose production is expanding and intensifying, and coffee production systems form a gradient of ecological complexity ranging from forest-like shaded polyculture to dense sun-exposed monoculture. We used LiDAR technology to survey the habitat, in combination with autonomous recording units and a vocalisation classification algorithm to assess bird community composition in a coffee landscape comprising a shade-grown coffee farm, a sun coffee farm, and a forest remnant, located in southern Mexico. We found that combining LiDAR with the automated analysis of continuously collected bioacoustics data can capture the expected functional signatures of avian communities as a function of habitat quality in the coffee landscape. Thus, we show that this approach can be a robust way to monitor how biodiversity responds to land use intensification in the tropics. A major advantage of this approach is that it has the potential to be deployed cost-effectively at large scales to help design and certify biodiversity-friendly productive landscapes.

## Introduction

Land use change is a leading driver of biodiversity loss worldwide (IPBES, 2019). In particular, the intensification of human land use is driving rapid declines in functional diversity, particularly in the tropics (Etard et al., 2022). Tropical forests, which are the most biodiverse terrestrial ecosystems on Earth, are threatened by an increasing demand for agricultural commodities (Laurance et al., 2014; Lewis et al., 2015), leading to an urgent need for policies that sustain both agricultural production and biodiversity in tropical regions. To stem the decline in tropical biodiversity, stakeholders engaged in supply chains must commit to sustainable agricultural practices (Rueda et al., 2017; Curtis et al., 2018), which rely on being able to quantify and monitor the response of biodiversity to habitat degradation.

An important crop produced in tropical forested regions is coffee. It is one of the world’s most traded commodities, being grown on over 1 million km^2^ of land worldwide (FAO, 2022), primarily located in highly biodiverse tropical mountains. Traditionally, coffee is grown under diverse and dense shade canopies, and it can have a vegetation structure and a level of biodiversity that is comparable to native forests (Perfecto et al., 1996; Alvarez-Alvarez et al., 2020). Therefore, traditional shade-grown coffee offers both a refuge for biodiversity in areas that have been hard-hit by deforestation (Perfecto et al., 1996) and a high-quality matrix connecting patches of protected forest (Perfecto & Vandermeer, 2010; Estrada-Carmona et al., 2022). However, a variety of socio-economic factors is leading to the expansion and intensification of coffee production across the world (Jha et al., 2014). Some of the main ways in which coffee production systems are intensified to increase yield include increasing sun exposure by a reduction in tree cover and applying chemical pesticides (Perfecto et al., 1996). Thus, coffee production systems now form a continuum ranging from forest-like shaded polyculture to dense sun-exposed monoculture, with an associated gradient of ecological complexity that provides a model system for investigating the response of biodiversity to habitat degradation (Perfecto et al. 2014).

To evaluate the habitat-biodiversity relationship in coffee and other tropical agricultural commodities, birds are commonly used as indicators as they are the best-known taxon, they occupy a wide range of ecological niches, and they are variably susceptible to disturbance (Sekercioglu et al., 2019). Birds are declining rapidly worldwide, and recent studies in temperate regions found that migratory birds that are spending their non-breeding season in the tropics are declining faster than temperate residents (Rosenberg et al., 2019, Burns et al., 2021). This decline potentially indicates associated widespread environmental degradation and biodiversity decline in tropical regions. Land use change is one of the leading drivers of bird decline (Bairlein, 2016; Sekercioglu et al., 2019) as it is associated with strong reductions in resource supply and in the availability of ecological niches. Consistently and reliably monitoring how bird communities are affected by land use change throughout the annual cycle in a way that can be scalable is important for certifying and examining the success of biodiversity-friendly agricultural practices, but it remains challenging using traditional survey methods.

Field surveys in coffee farms have been conducted by human observers using point counts and mist-netting methods to evaluate avian diversity and using visual methods to estimate foliage structure (Greenberg et al., 1997; Tejeda-Cruz & Sutherland, 2004; Sekercioglu et al., 2019; Ong’ondo et al., 2022). These approaches, however, are labour intensive and with limited reproducibility, which affects their scalability for widespread use. In this study, we examined whether the automated analysis of audio data collected by passive acoustic monitoring (PAM), together with remote sensing, can help with monitoring avian biodiversity along the gradient of habitat degradation associated with the intensification of coffee production. Specifically, we used autonomous recording units (ARUs) and a vocalization classification algorithm to assess bird community composition, in combination with Light Detection And Ranging (LiDAR) technology to quantify habitat quality, in a coffee landscape comprising a shade-grown coffee farm, a sun coffee farm, and a forest remnant. Automated acoustic data processing was used here to monitor avian community composition continuously throughout a full annual cycle, which enabled the examination of community change between seasons with different climate (e.g., dry versus rainy), and the associated periods of the year when long-distance migratory species are present, passing through, or largely absent. This full annual cycle perspective is crucial for understanding the factors that are structuring bird communities (Herrera, 1978) and how they respond to habitat change.

To evaluate how well bird diversity and habitat can be monitored in a consistent and scalable way across coffee landscapes, we investigated whether the combination of bioacoustics and LiDAR can capture general patterns in avian communities and vegetation that are regularly observed along tropical land use intensity gradients. We tested the hypothesis (H1) that LiDAR can capture the association between the intensification of coffee production and a corresponding decrease in vegetation density and complexity (Perfecto et al., 1996; Moguel & Toledo, 1999). We also tested the following three hypotheses related to how bioacoustics can capture the response of functional signatures of avian communities to coffee production and its intensification: (H2) the proportion of forest specialists is higher in the forest while the proportion of generalist species increases with habitat disturbance (Greenberg et al., 1997; Tejeda-Cruz & Sutherland, 2004; Newbold et al., 2013; Buechley et al., 2015; Sekercioglu et al., 2019; Jarrett et al., 2020; Ong’ondo et al., 2022); (H3) the proportion of migrant species, which tend to be generalist and opportunistic, increases with habitat degradation (Greenberg et al., 1997; Tejeda-Cruz & Sutherland, 2004; Newbold et al., 2013; Buechley et al., 2015); and (H4) the proportion of invertivores and understory specialists decreases with habitat degradation (Newbold et al., 2013; Buechley et al., 2015; Sekercioglu et al., 2019; Jarrett et al., 2020).

## Materials & Methods

### Study Site

Data was collected between February 2022 and February 2023 in a coffee landscape located between 900-1300m a.s.l. in the Soconusco region of Chiapas in southern Mexico. Most of the native forest in mountainous parts of the region has been converted for coffee agriculture (Philpott et al., 2008), making it a coffee-growing agricultural matrix. However, the terrain is rugged (Fig. 1b) and farmers often leave forest patches on steep banks. The coffee landscape is composed of (i) a 278-ha shade-grown coffee farm (Finca Irlanda, Fig. 1d), with canopy cover comprised primarily of nitrogen-fixing *Inga* species, though up to 200 species of tree have been recorded on the farm which is classified as ‘diverse shade’ (Perfecto *et al*. 2003); (ii) a 300-ha sun coffee farm (Finca Hamburgo, Fig. 1e), where canopy cover and vegetation diversity are significantly lower than in Irlanda, although it is interspersed with patches of moderate shade, and which makes use of synthetic agrochemicals and as such is classified as a ‘low diversity shaded non-organic coffee farm’ (Perfecto et al., 2003); and (iii) a forest fragment composed of primary and secondary forest and located between the two farms (Fig. 1c). Autonomous acoustic monitoring units (Fig. 1f) were deployed, and LiDAR data were collected, in each zone of this coffee landscape (i.e., shade coffee, sun coffee, and forest fragment).

**Figure 1:**
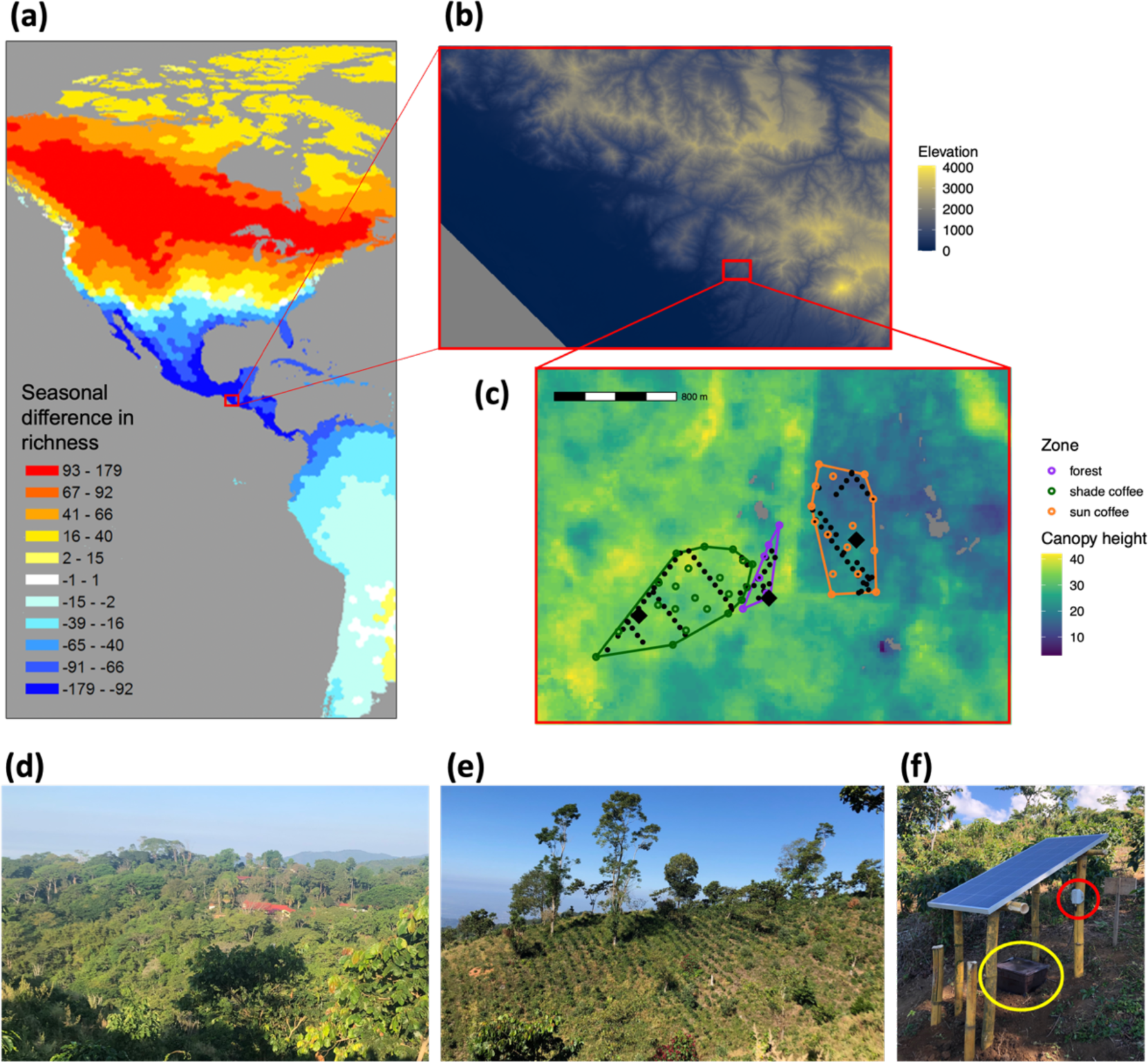
Study site. (a) Map showing the pattern of seasonal difference in avian richness across the Americas, calculated as the number of species present only during their breeding season minus the number of species present only during their non-breeding season (taken from Somveille et al., 2013). The red rectangle indicates the location of the study region on the Pacific slope of Chiapas, Mexico, whose topography is shown in (b). (c) Canopy height map obtained from Lang *et al*. (2023), covering the study area which includes a shade coffee farm (Finca Irlanda, in bottom-left quadrant), a sun coffee farm (Finca Hamburgo, in top-right quadrant), and forest patches. Open circles indicate the location of LiDAR scans obtained using the hand-held device; polygons indicate minimum convex hulls around the hand-held LiDAR scans for each zone; Closed points indicate the location of GEDI footprints falling within each polygon; Black diamonds indicate the location of the autonomous acoustic monitoring units that were deployed in sun coffee, shade coffee and forest. Photos show (d) the shade coffee farm, (e) the sun coffee farm, and (f) an autonomous acoustic monitoring unit composed of a solar panel, a battery (in yellow circle) and a BUGG acoustic recording device (in red circle).

### Habitat structure

LiDAR is an active remote sensing technology that emits laser pulses and measures the travel time of the emitted pulses i.e. their distance and position in three dimensions. LiDAR has been widely used to measure the three-dimensional structure of forests (Yao et al., 2012; Asner et al., 2014). We used two types of LiDAR data: (i) scans collected from the ground using a handheld device, and (ii) scans collected from space from a device orbiting the Earth. These two LiDAR datasets were then compared to examine their capacity to capture the difference in vegetation between shade coffee, sun coffee, and forest.

### Handheld LiDAR data collection

Vegetation data was collected at several locations within each zone of the coffee landscape using a GeoSLAM ZEB Revo RT handheld LiDAR scanner (hereafter referred to as handheld LiDAR). Operation of the device followed the instructions set out in the user manual. Scans were centred on a given point, from which the operator walked multiple overlapping figures-of-eight, following the procedure set out by Bauwens et al. (2016), and overall bounded at a radius of 25m for a sampling duration of 5-7 minutes, depending on the accessibility of the terrain. Each plot where an autonomous acoustic monitoring device was deployed was scanned using the handheld lidar device. Several additional plots were also sampled randomly across each of the different areas in the coffee agroecosystem, according to the size of the area, and each separated by a minimum distance of 200m: 8 plots in forest, 15 plots in sun coffee and 22 plots in shade-grown coffee (Fig. 1c). At every location where vegetation data was collected, we visually recorded an estimate of canopy cover as the proportion of sky obstructed by vegetation as seen from the ground.

### Handheld LiDAR data pre-processing

Point clouds obtained by the handheld LiDAR scans were first classified into ground and non-ground points using the Cloth Simulation Function algorithm (Zhang et al. 2016). Point clouds were height-normalised using digital terrain models (DTMs) based on spatial interpolation using a k-nearest neighbour (with k=10) approach with an inverse-distance weighting (with power p=2). DTMs model the ground points of a point cloud, which were then subtracted from the original point cloud, resulting in a height-normalised point clouds in which topography is smoothed such that metrics pertaining to vertical stratigraphy are not confounded by the slope of terrain. Ground points were removed after the height-normalisation so that the remaining points solely represented vegetation. Finally, point clouds were clipped at a radius of 25m from their origin, to remove outlying points after the point at which LiDAR returns diminish due to occlusion. All the data pre-processing was conducted using the R packages lidR (Roussel et al. 2020).

### Handheld LiDAR vegetation metrics

We first projected a three-dimensional voxel grid onto the height-normalised point clouds to divide them into 1 m^3^ voxels and obtain Leaf Area Density (LAD) values per voxel. The voxel area of 1 m^3^ was chosen as a compromise of maximising resolution while avoiding high computational time. A vertical LAD profile (in m^2^/m^3^) was calculated for each handheld LiDAR scan, which gives the vertical distribution of one-sided leaf surface area per layer of a given height, across a given horizontal area. Based on the LAD profile, the Leaf Area Index (LAI; in m^2^/m^2^), which measures one-sided leaf surface area per unit of ground surface (Watson, 1947), was calculated for understory (0-5m), mid-height vegetation (5–10m), canopy (10-20m) and high canopy (>20m). We also calculated the Foliage Height Diversity (FHD), which is a measure of the complexity of vegetation structure that has been shown to be a good indicator of habitat quality for birds and to correlate with bird diversity (MacArthur & MacArthur, 1961), as well as the maximum canopy height as the maximum height for which LAD > 0.001 across the LAD profile. All vegetation metrics for the handheld LiDAR scans were calculated using the R package leafR (methodology presented in de Almeida et al., 2019).

### Spaceborne LiDAR data

Spaceborne LiDAR data was obtained from NASA’s Global Ecosystem Dynamic Investigation (GEDI) system, which provides high-resolution laser ranging using an instrument mounted on the International Space Station that samples footprints of 25m diameter every 60m, with a pointing uncertainty of 10m (Dubayah et al., 2020). We used GEDI Level 2B data products (Dubayah et al., 2021) between February 2019 and February 2024. GEDI footprints for each zone (i.e. shade coffee, sun coffee, and forest fragment) were extracted within polygons representing convex hulls calculated using the geographical locations of the handheld LiDAR scans (Fig. 1c). A total of 36 footprints were obtained in the sun coffee zone, 39 footprints were obtained in the shade coffee zone, and 13 footprints were obtained in the forest zone. For each GEDI footprint, Plant Area Index (PAI, in m^2^/m^2^) was calculated for understory (0-5m), mid-height vegetation (5–10m), canopy (10-20m), and high canopy (>20m). PAI is highly comparable to LAI for describing vegetation structure across our landscape, as both measures have been shown to be closely related (Ziegler et al., 2023). In addition, as the point clouds from the handheld LiDAR scans did not differentiate between green and non-green components of the vegetation, thus LAI (calculated for handheld LiDAR) is directly comparable to PAI (calculated for spaceborne LiDAR). We also obtained the FHD as well as the maximum canopy height calculated as the height above ground of the received waveform signal start. All the GEDI data processing was conducted using the R package rGEDI.

### Avian community composition and dynamics

#### Data collection

To monitor the composition of avian communities across the coffee landscape, audio data were collected via passive acoustic monitoring using recording devices deployed at three locations: in the shade coffee zone (15°10’02.7°N, 92°20’37.4°W), in the sun coffee zone (15° 10’ 18.696°N, 92’ 19’ 49.584’W), and in the forest fragment (15’ 10’ 6.384’N, 92’ 20’ 8.808’W); Fig. 1c). We used solar-powered BUGG acoustic recording devices (Sethi et al., 2017; www.bugg.xyz; Fig. 1f), which collected audio data between 20 February 2022 and 20 February 2023 using Knowles SPH0641LU4H-1 MEMS microphones (sensitivity -26dB @ 94dB SPL). At each site, audio data was recorded continuously 24 hours a day at a sampling frequency of 44.1 kHz, except when the battery could not be sufficiently charged by the solar panel due to cloud and vegetation cover, and then saved into 5-min audio files that were compressed using FFMPEG’s implementation of LAME VBR0 MP3 compression. The audio data was retrieved once a month from the devices to be analysed.

#### Data processing for species detections

Automatic vocalisation detection was conducted on the collected audio data using BirdNET, the largest open-source bird vocalisation classification neural network model available (Kahl et al., 2021), which can identify over 6000 bird species and is highly precise in many regions of the world (Sethi et al., 2023). We used the BirdNET-Analyzer model v2.3 and location data was provided to BirdNET to filter for only species expected to occur in the region based on eBird observations.

To calibrate BirdNET, for each species, we randomly selected 50 detections with BirdNET classification confidences over 0.80. Species with fewer than 50 detections across all sites and throughout the recording period were discounted, thus retaining 59 species overall. We listened to the randomly selected detections and labelled them as correct, incorrect, or unsure (e.g., if the call was ambiguous). Conservatively, we denoted unsure labels as false positives.

BirdNET precision, *p*, was measured as *p = T_p_/T_p_+F_p_*, where *T_p_* and *F_p_* denote true and false positives, respectively. For the rest of the analyses, we only considered species with over 80% precision, which corresponds to 50 out of 59 species.

#### Species’ temporal ranges

For each site (i.e. acoustic detections in shade coffee, sun coffee, and forest), we only considered species with more than 20 detections over the whole annual cycle in that site. This filter was applied in order to have a sufficient number of detections per species in order to estimate their temporal ranges in a given site. The species detection datasets (one per site) were split into 73 5-day periods between 20 February 2022 to 20 February 2023. Adapting a methodology that has been used for analysing species detections in eBird checklists (Freeman et al., 2022), for each period and each species, we calculated a detection rate as the number of 5-min audio files with a detection for this species divided by the total number of 5-min audio files, which was then rescaled between 0 and 1 by dividing by the maximum detection rate value. The species was considered present during a given period if the detection rate was above 0.05. To limit the effect of temporal variation in recording effort and variation in species detectability, which can generate false absences (i.e. period where the species’ detection rate is not above 0.05 despite the species being locally present during that period), we assumed that if the species was absent for fewer than eight 5-day periods (40 days) between two periods of presence, then the species was considered present continuously during that time span. This assumes that the species must be absent for at least 40 days to be considered as having migrated away from the area (and back), otherwise, it is considered as being present but not detected. We also considered that a period of presence that is separated from any other period of presence by more than four 5-day periods (20 days) is a false presence and therefore was set to an absence.

#### Functional signatures of avian assemblages

We obtained the following functional trait data for bird species from the AVONET dataset (Tobias et al., 2022): migration, habitat, habitat density, and trophic niche. In addition, we collected data on foraging strata for the species detected in our dataset from Birds of the World (Billerman et al., 2022), which we encoded as binary values for foraging in three strata: understory, mid-height vegetation and canopy (a species can forage in more than one strata). From these trait data, we calculated the following functional signatures of avian assemblages for each 5-day period at each site:

*Proportion of forest specialists*: number of occurring species whose habitat is classified as “forest” and habitat density classified as “dense habitat”, divided by the total number of occurring species. Upon inspection of the forest specialist species, the following species were not classified as forest specialists despite meeting the habitat and habitat density criteria as they commonly occur in non-forested habitats across their range: Wilson’s Warbler, Magnolia Warbler, Black-and-white Warbler, Western Tanager, Gray Hawk, Roadside Hawk, Rufous-tailed Hummingbird.

*Proportion of understory specialists*: number of occurring species that are foraging only in understory divided by the total number of occurring species.

*Proportion of strata generalists*: number of occurring species that are foraging on all three strata (understory, mid-height vegetation and canopy) divided by the total number of occurring species.

*Proportion of migrants*: number of occurring species that are seasonal migrants divided by the total number of occurring species.

*Proportion of invertivores*: number of occurring species that forage on invertebrates divided by the total number of occurring species.

*Proportion of omnivores*: number of occurring omnivorous species divided by the total number of occurring species.

#### Sensitivity analysis

To assess whether the resulting functional signatures of avian assemblages are sensitive to our choice of parameter values used for data processing for species detections and temporal ranges, we conducted a sensitivity analysis on the following parameters: BirdNET precision threshold (set value: 0.8; we also ran the analysis for values of 0.7 and 0.9), minimum number of species detections over the whole annual cycle at a site (set value: 20; we also ran the analysis for values of 10 and 30), false absence maximum time span (set value: 40 days; we also ran the analysis for values of 20 and 60 days), and false presence maximum separation period (set value: 20 days; we also ran the analysis for values of 10 and 40 days). We ran the analysis by changing the value of only one parameter at a time and generated the functional signatures described above.

## Results

Vegetation scans from the handheld LiDAR device indicated a decrease in foliage density, as measured by LAI, with increasing vegetation height for all three study zones (Fig. 2a). No significant differences in LAI above the understory were detected between all three study zones, while in the understory, LAI was significantly higher in the forest fragment compared to sun coffee (pairwise t-test, P_adj_=0.01) and shade coffee (pairwise t-test, P_adj_=0.021). Handheld LiDAR scans at the sites of deployment of the acoustic monitoring units showed similar LAI trends as the scans across the study zones but tended to have higher LAI values than the rest of the shade coffee scans above 5m (Fig. 2a).

**Figure 2:**
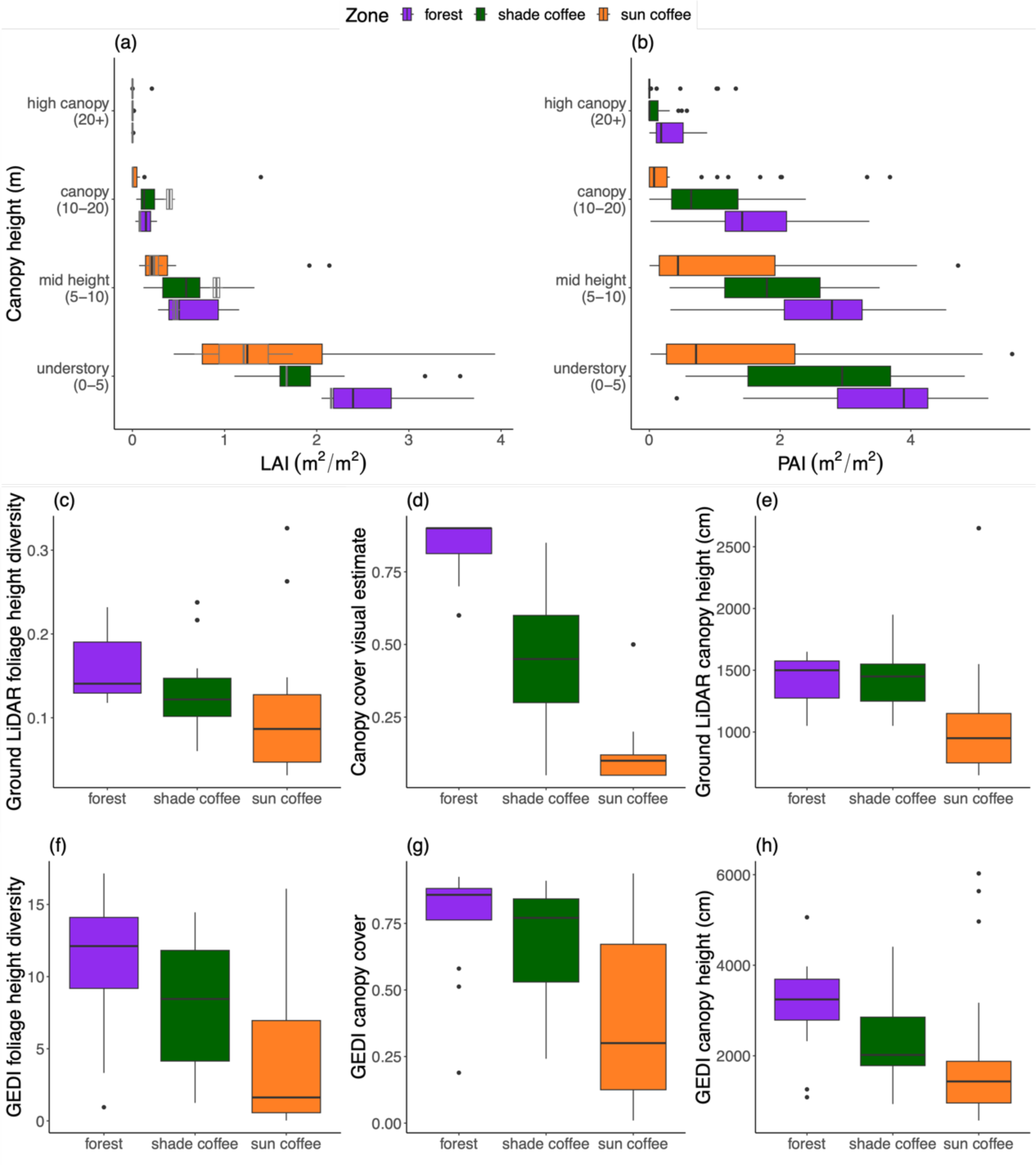
Vegetation structure measured by LiDAR. (a) Leaf Area Index (LAI) estimated using the hand-held device for different ranges of height for each zone; grey boxplots indicate the values obtained at the exact location of deployment of the autonomous acoustic monitoring units at the start and end of the year. (b) Plant Area Index (PAI) estimated using GEDI footprints for different ranges of height for each zone. Foliage Height Diversity (FHD) is shown for each zone as calculated based on (c) ground handheld LiDAR and (f) GEDI. Canopy cover is shown for each zone as calculated based on (d) visual estimate and (g) GEDI. Canopy height is shown for each zone as calculated based on (e) ground handheld LiDAR and (h) GEDI. The boxplots indicate the first, second, and third quartiles of the variation in vegetation metrics across the hand-held LiDAR scans and GEDI footprints obtained within each study zone (Fig. 1c).

In agreement with the handheld LiDAR scans, spaceborne LiDAR scans from GEDI indicated a decrease in foliage density, as measured by PAI, with increasing vegetation height for all three study zones (Fig. 2b), but this decrease was less abrupt and more linear. However, in contrast with handheld LiDAR, spaceborne LiDAR indicated that PAI in sun coffee was consistently significantly lower than in forest (pairwise t-tests – understory: P_adj_=0.004, mid-height vegetation: P_adj_=0.004, canopy: P_adj_=0.013; Fig. 2b), and significantly lower than in shade coffee below 10m (pairwise t-tests – understory: P_adj_=0.016, mid-height vegetation: P_adj_=0.017, canopy:

P_adj_=0.167; Fig. 2b). Spaceborne LiDAR also indicated that PAI is not significantly different in forest than in shade coffee across all vegetation strata (pairwise t-tests – understory: P_adj_=0.393, mid-height vegetation: P_adj_=0.230, canopy: P_adj_=0.136; Fig. 2c).

The response of vegetation to the land use intensity gradient in the coffee agroecosystem exhibited similar trends for FHD, canopy cover and canopy height as measured by spaceborne LiDAR, with forest tending to have higher values than the coffee plots, and shade coffee having higher values than sun coffee (Fig. 2f-h). These results largely matched the ones obtained from the ground using handheld LiDAR and visual canopy cover estimates (Fig. 2c-e), except for canopy height that was not higher in the forest than in shade coffee based on handheld LiDAR scans (Fig. 2e).

The autonomous acoustic monitoring units recorded 17,654 hours of audio data together throughout the study period. Whilst recording effort was consistent and nearly continuous in both shade and sun coffee (despite a couple of drops in sun coffee), it was significantly lower and varied substantially in forest, with a broad decrease during the rainy season (May–September; Fig. 3a). Overall, applying BirdNET to our audio data yielded 123,355 species detections throughout the study period. The total number of species detections was substantially higher in shade and sun coffee than in forest during the rainy season, but it was more variable during the dry season (Fig. 3b). Detection rate (number of detections per minute), however, was more comparable between the three sites (Fig. 3c).

**Figure 3:**
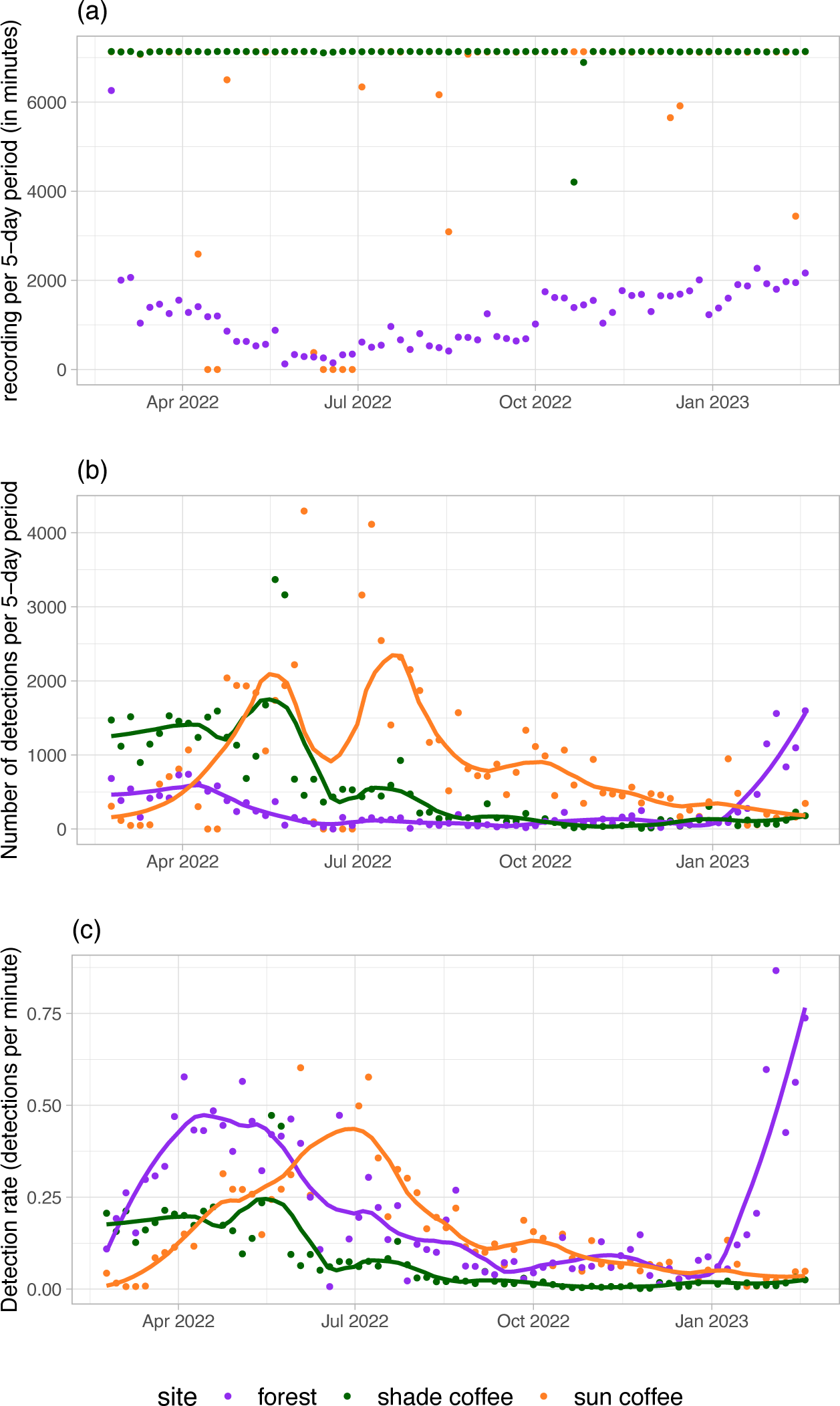
Acoustic recording effort. Recording effort (a), detections (b) and detection rate (c) per 5-day period collected by each acoustic monitoring unit throughout a full annual cycle. Lines indicate smoothing curves fitted using local polynomial regression with a smoothing parameter of α = 0.25.

Overall, we were able to estimate the temporal ranges of occurrence of 50 species across our 3 sites and throughout the study period (Fig. 4). In sun coffee, 32 different species were recorded for a cumulated 7105 days of presence (sum across all species of days when a given species is considered present). This was lower than in shade coffee, where 45 different species were recorded for a cumulated 9220 days of presence. Finally, in forest, 26 different species were recorded for a cumulated 5210 days of presence. However, because of the significantly lower recording effort in the forest (Fig. 3), alpha diversity measures could not be straightforwardly compared to the coffee plots.

**Figure 4:**
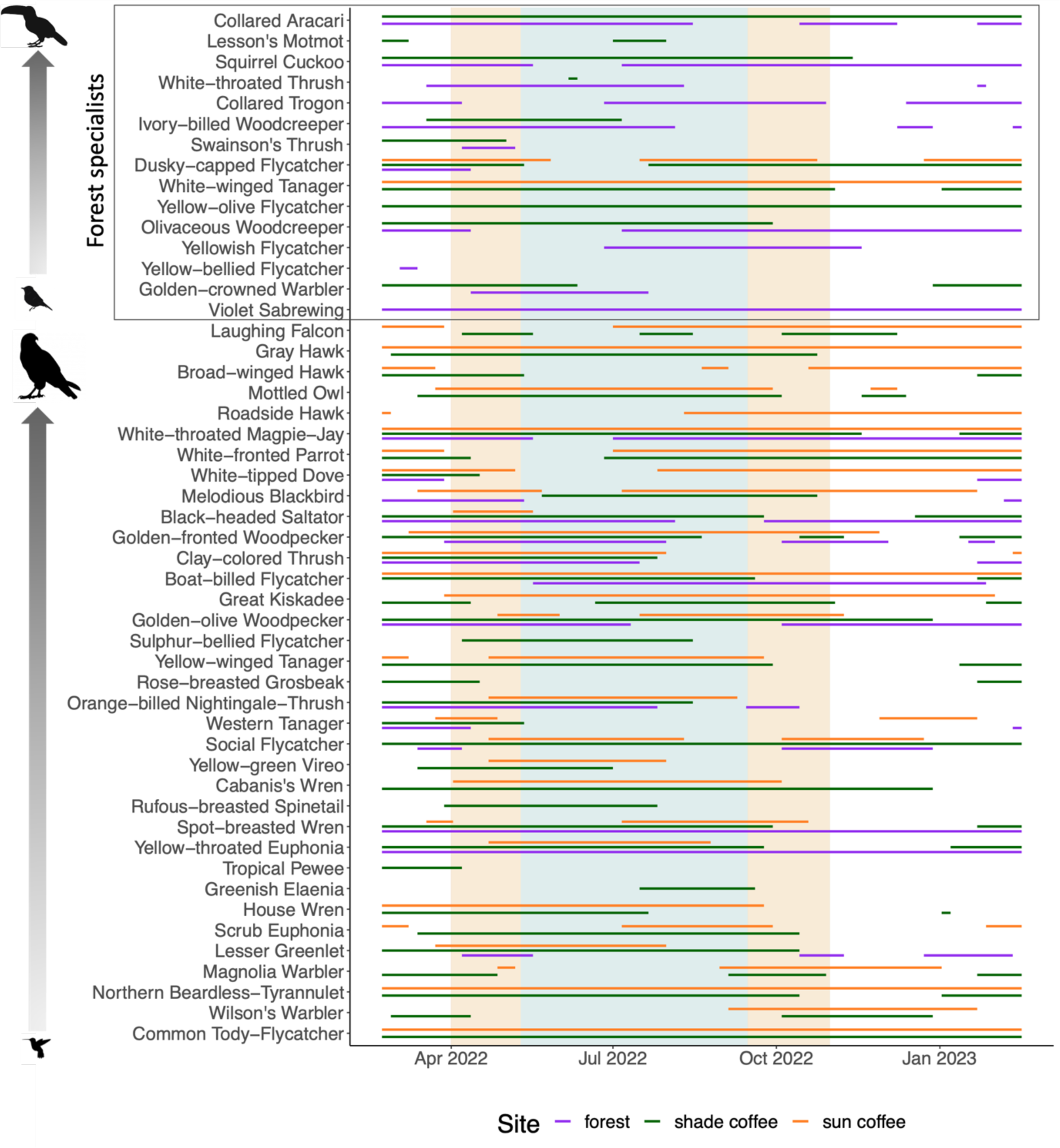
Species’ temporal ranges in the study area. Lines indicate the range of presence of a species in a given site (shade coffee, sun coffee, or forest). Species are ordered in descending order by body mass, with forest specialists separated from the rest. The blue vertical range in the background indicates the rain season, and the orange vertical ranges indicate the shoulder seasons between rain and dry seasons.

The proportion of forest specialists was consistently higher in the forest than in the coffee plots throughout the year, and consistently higher in shade coffee than in sun coffee (Fig. 5a,g). In the forest, the proportion of forest specialists was significantly higher during the rainy season (two-sample t-test: t=-7.397, P<0.001), while in shade coffee it was significantly higher during the dry season (t=5.868, P<0.001; Fig. 5g). The proportion of understory specialists is nearly consistently higher in forest compared to coffee plots, particularly during the dry season, and consistently higher in sun coffee than in shade coffee during the dry season (but not during the rainy season; Fig. 5h). The proportion of strata generalists varies greatly throughout the study period for all three sites without evident patterns associated with seasonality, but it is nearly consistently higher in sun coffee than in the forest during the rainy season, with shade coffee somewhat in between (Fig. 5c). In the forest and shade coffee, the proportion of strata generalists was significantly lower during the dry season compared to the rainy season (forest: t=4.206, P<0.001; shade coffee: t=3.157, P=0.003) while no seasonal difference is observed in sun coffee. The proportion of migrants is particularly low in the forest, and consistently lower than in either coffee sites (Fig. 5d). This proportion is similar in both coffee sites during the rainy season, but it exhibits wide variations during the dry season, with the first part of the dry season (November–December) corresponding to a peak in the proportion of migrants in sun coffee and a dip in shade coffee. The later, drier part of the dry season (February–March) corresponds to a peak in the proportion of migrants in shade coffee as well as in the forest (Fig. 5d). The proportion of invertivores is consistently higher in the forest than in sun coffee, while it is similar between sun and shade coffee during the rainy season but consistently higher in shade coffee during the dry season (Fig. 5e). In sun coffee, the proportion of invertivores is significantly lower during the dry season compared to the rainy season (t=-8.130, P<0.001), while the opposite is observed in shade coffee (t=2.794, P=0.008). Finally, the proportion of omnivores is nearly consistently higher in sun and shade coffee than in forest, without much seasonal variation except for peaks in forest and shade coffee during spring migration and late dry season, respectively, and drops in forest and shade coffee during late rainy season and early dry season respectively (Fig. 5f).

**Figure 5:**
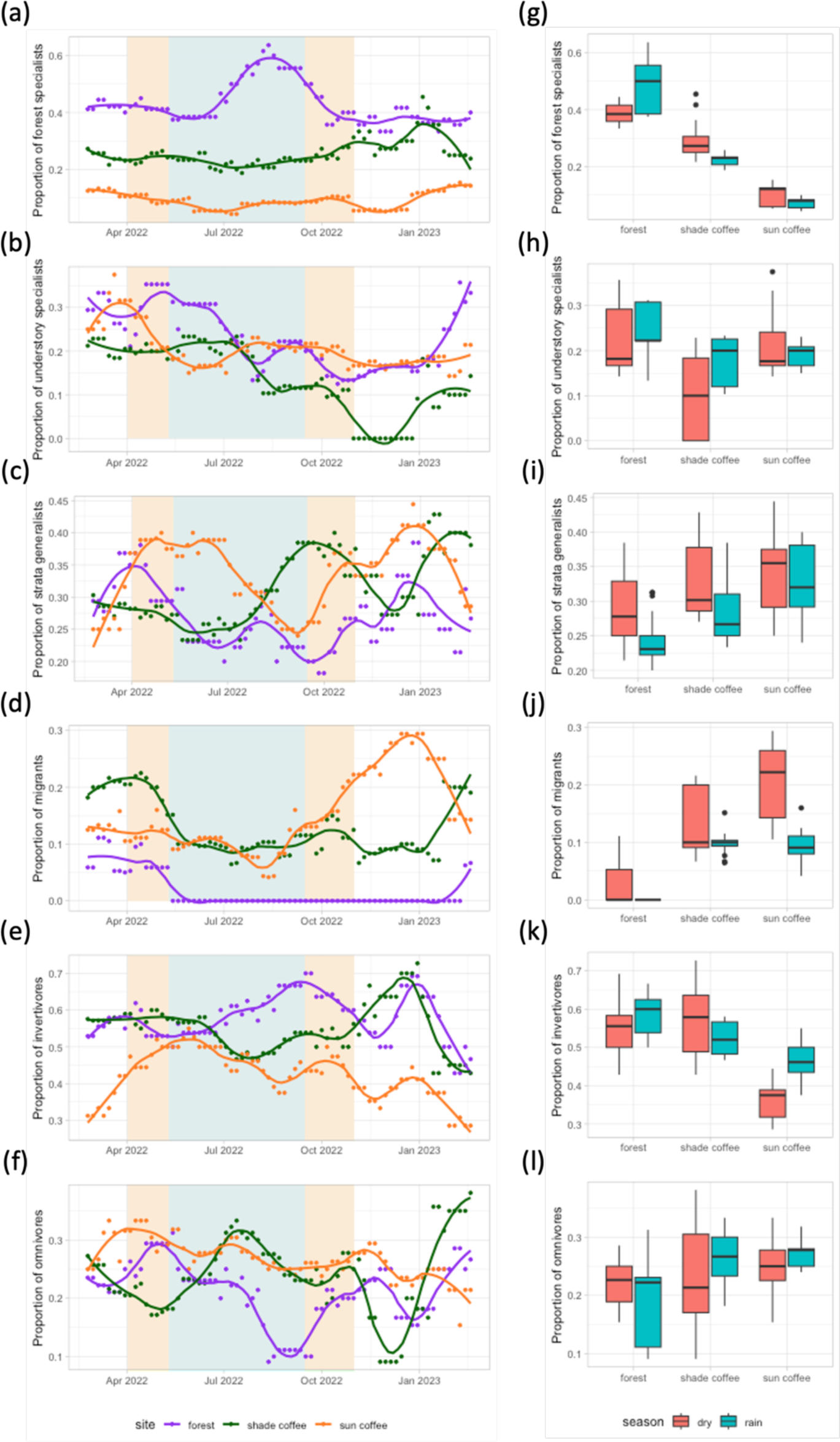
Annual trends in functional signatures. Functional signatures per 5-day period throughout the annual cycle (a–f) and per season (g–l) for each site (shade coffee, sun coffee, and forest), showing trends in the proportion of (a, g) forest specialists, (b, h) understory specialists, (c, i) strata generalists, (d, j) migrants, (e, k) invertivores, and (f, l) omnivores. Lines indicate smoothing curves fitted using local polynomial regression with a smoothing parameter of α = 0.25.Figures S1–S8 show the results of the sensitivity analysis, which assess how the functional signatures of avian communities captured through bioacoustics vary with changes in parameter values used for data processing for species detections and temporal ranges.

The sensitivity analysis indicated that varying the parameter values did not substantially affect the results. The variation in functional signatures of avian communities between seasons and sites remained qualitatively similar for all combinations of parameter values (Figs. S1–S8), thus our conclusions are not sensitive to specific choices of parameter values.

## Discussion

Monitoring how habitat quality and animal communities respond to land use change is critical for certifying and examining the success of biodiversity-friendly agricultural practices, particularly in the tropics. Our results demonstrate that the automated analysis of audio collected by PAM, together with the processing of remote sensing data, can be used to monitor bird communities and vegetation structure consistently and reliably along a tropical land-use intensity gradient. Firstly, we find that LiDAR technology can capture expected habitat quality metrics in shade-grown coffee, sun coffee, and forest remnants. Secondly, we find that integrating ARUs with the BirdNET vocalization classification algorithm can capture the expected functional signatures of avian communities as a function of habitat quality in a coffee agroecosystem.

Coffee production and its intensification are associated with a corresponding decrease in vegetation density and complexity which ultimately affects habitat quality for animals (Perfecto et al., 1996; Moguel & Toledo, 1999). Our results indicate that LiDAR technology can capture this pattern. Quantitative metrics derived from LiDAR scans to estimate vegetation density and complexity, namely LAI/PAI, FDH, canopy cover, and canopy height, are all markedly higher in the forest than in sun coffee, with shade coffee in between the two (Fig. 2), thus supporting hypothesis H1. We show that GEDI can reliably capture the habitat degradation gradient in the coffee agroecosystem, qualitatively matching estimates obtained from the ground. Handheld LiDAR is able to estimate vegetation structure in the understory but appears limited for estimating vegetation density and structure in the canopy as it cannot differentiate between the different sites above the understory for both LAI and canopy height (Fig. 2). Our results, therefore, suggest that GEDI data, which is freely available globally and does not require labour on the ground, is potentially a powerful and appropriate tool for estimating habitat quality along gradients of human land-use intensity in tropical forests. It must be noted, however, that additional information on the richness and composition of tree species present in shade coffee and forest, and on the use of chemical pesticides, will be important to fully assess how habitat shapes animal biodiversity (Philpott et al., 2008; Perfecto & Vandermeer, 2008; Bakermans et al., 2012; Gebremichael et al., 2022). In addition, a limitation of GEDI data that could apply to our study is that the vertical resolution of its footprints can be reduced in mountainous terrain as the canopy signal in a 25m footprint is affected by steep slope (Fayad et al., 2021). As we used many footprints broadly distributed across the patches of coffee and forest (Fig. 1c), the broad comparison of vegetation structure between sites might not be significantly affected by this issue. However, as patches of forest remnant tend to be located on steeper slopes in coffee growing regions, this limitation will necessitate further examination.

Birds are excellent indicators of ecosystem health as they occupy a wide range of ecological niches, and they are affected by the intensification of coffee production (Greenberg et al., 1997; Tejeda-Cruz & Sutherland, 2004; Sekercioglu et al., 2019). Our results indicate that the automated analysis of audio collected by PAM can capture known patterns in how coffee production and its intensification affect various functional signatures of avian communities. Firstly, in support of hypothesis H2, we found that the proportion of forest specialists is higher in forest (Fig. 5a,g) while the proportion of generalists (i.e. strata generalists and omnivores) tends to increase with increasing habitat degradation associated with the intensification of coffee production (Fig. 5c,i,f,l; Greenberg et al., 1997; Tejeda-Cruz & Sutherland, 2004; Newbold et al., 2013; Buechley et al., 2015; Sekercioglu et al., 2019; Jarrett et al., 2020; Ong’ondo et al., 2022). Secondly, in support of hypothesis H3, we found that the proportion of migrant species increases with increasing habitat degradation (Fig. 5d,j; Greenberg et al., 1997; Tejeda-Cruz & Sutherland, 2004; Newbold et al., 2013; Buechley et al., 2015). Finally, in support of hypothesis H4, we found that the proportion of invertivores decreases with increasing habitat degradation (Fig. 5e,k; Newbold et al., 2013; Sekercioglu et al., 2019; Jarrett et al., 2020). We also found that the proportion of understory specialists tends to be higher in forest than in coffee, in agreement with hypothesis H4 (Buechley et al., 2015; Sekercioglu et al., 2019; Jarrett et al., 2020). However, during the dry season this proportion tends to be higher in sun coffee than in shade coffee (Fig. 5b,h), which warrants further investigations to determine whether this pattern is generally observed across coffee agroecosystems. Overall, our results support the use of bioacoustics as a robust tool for estimating avian community composition along gradients of human land-use intensity in tropical forests. Passive acoustic monitoring based on ARUs is more and more widespread in ecological research (Shonfield & Bayne, 2017; Ross et al., 2023) and increasingly affordable. Together with the availability of vocalization classification algorithms like BirdNET to detect species from the recorded audio data, it renders the use of bioacoustics much more accessible and scalable (Wood et al., 2022; Sethi et al., 2023). We show here that PAM can be leveraged for biodiversity conservation in coffee landscapes.

In this study, we used an automated approach to processing bioacoustics data based on power-autonomous devices that continuously record audio data to be analysed by BirdNET. This approach allows for the consistent collection of data throughout the full annual cycle, which has previously not been possible with traditional survey methods, and it can thus contribute to a better understanding of how habitat degradation affects monthly and seasonal changes in community structure. In particular, our automated approach is very pertinent in mid-elevation areas in the northern tropics, where much of the world’s coffee is produced, and which are home to highly dynamic bird communities that host many non-breeding migrants during the northern winter (Fig. 1a; Somveille et al., 2013), many passage migrants during migration, and altitudinal migrants. These regions are therefore well suited for examining the effect of land use intensity on the seasonal cycle of avian communities. Our results reveal that while several functional signatures of avian communities remain relatively stable throughout the year, some do vary. Firstly, the proportion of migrants tends to be higher during the dry season across the study area, which makes sense given the geographical location in a region of high richness in non-breeding migratory birds from the northern hemisphere (Fig. 1a). Secondly, in the forest during the rainy season, which corresponds to the breeding season for most species in southern Mexico, the proportion of forest specialists is higher than for the rest of the year (Fig. 5a,g) and the proportion of strata generalists is lower (Fig. 5c,i). In sun coffee, the proportion of invertivores is lower during the dry season than during the rainy season (Fig. 5e,k). These results indicate that forest specialists are potentially particularly dependent on the forest during the rainy, breeding season. It also suggests that diet specialists, particularly those relying on forest resources (i.e. invertivores), are more affected by habitat degradation during the dry season.

In addition to the observed seasonal variation in functional signatures of avian communities between the dry and rainy seasons, our results also reveal intriguing temporal patterns within the dry season that, to our knowledge, have not been previously described. Firstly, we found that habitat and diet generalists (i.e. strata generalists, omnivores, and migrants) tend to rely more on degraded habitat during the first part of the dry season and then rely more on shaded and forested habitat during the second part of the dry season. This pattern of change in habitat use within the dry season for generalists might be due to a worsening of environmental conditions in sun coffee as the dry season progresses, potentially associated with disturbance related to coffee harvesting, or a relative increase in the availability of resources or microclimatic conditions in shade coffee and forest in the late dry season. Corroborating previous findings (Bakermans et al., 2009), our results therefore suggest that shade coffee might be a refuge for generalists during the late dry season. Secondly, we found that the proportion of invertivores and the proportion of understory specialists appear to respectively decrease and increase in all three study zones as the dry season progresses (Fig. 5). This pattern could be driven by altitudinal migration as birds might move along the elevational gradient during the dry season to track favourable conditions. Changes in the structure and density of vegetation might also occur throughout the dry season and could potentially drive some of the observed patterns. However, investigating this hypothesis requires obtaining enough LiDAR data at high spatio-temporal resolution, which is currently not available from GEDI. Finally, it is worth noting that the dry season of the year during which we monitored the coffee landscape was particularly dry, which might explain why generalists use shade coffee and forest as a refuge and why invertivores decreased as the dry season progressed. Climate change might result in extended periods of drought in coffee growing regions, which would enhance the conservation value of shade coffee and forest fragments as a refuge for birds.

While bioacoustics appears to be a robust tool for monitoring the functional signatures of avian communities along coffee production intensity gradients, it relies on several assumptions and has limitations. Firstly, bioacoustics technology is only able to detect vocal species, which might represent a limited subset of the local avian communities. Adding to this bias, bird vocalisation classification models like BirdNET necessitate large training datasets which tend to only be available for common North American and European species. Secondly, many forest birds tend to be more vocal and detectable during the breeding season, which may lead to seasonal biases in detections. We indeed found that detection rates were higher during the spring, which corresponds to the breeding season for most birds in our study region, with peaks in April-May in the forest and shade coffee and in May-July in sun coffee (Fig. 2c). Thirdly, the acoustic signal does not propagate uniformly depending on habitat structure (Ross *et al*. 2023), resulting in reductions in detection range as the vegetation gets denser. As we are investigating bird communities along a gradient of vegetation density and complexity, this might be associated with a corresponding gradient in detection probability. Finally, in addition to the increased occlusion of acoustic signals, the gradient in vegetation density and complexity is associated with a gradient in the occlusion of light, which leads to a lower capacity for recharging the devices’ batteries using solar power with increasing canopy cover. Recording effort was indeed markedly lower in forest, particularly during the rainy season (Fig. 2a), which results in a limited capacity for comparing our estimates of species temporal ranges between the different sites, thus affecting our ability to investigate alpha diversity across sites and throughout the year. Overall, as we focus on functional signatures of avian communities that are based on the proportional occurrence of functional groups (Sekercioglu et al., 2019), we assumed that these limitations do not generate biases in the acoustic detection probability for species from the different functional groups of interest, e.g. generalists versus specialists. More research is however needed to investigate whether this assumption is valid to support the use of bioacoustics for monitoring avian communities in coffee agroecosystems. More generally, more methodological development and extensive validation of acoustic monitoring technology are required to reduce the biases and limitations described above.

In this study, we show that combining remote sensing and the automated analysis of bioacoustics data can be a robust approach to monitoring how avian communities respond to land use intensity in the tropics. Specifically, spaceborne LiDAR can be used to assess the vegetation structure profile of a delineated patch of land and determine how degraded it is compared to tropical forest, and bioacoustics can be used on the ground to evaluate the state of biodiversity in that patch using the functional signatures of the avian community as indicators. To scale this methodology and be able to use it for testing the efficacy of sustainable agricultural practices and certifying biodiversity-friendly production of tropical agricultural commodities, several steps are needed for future work. Firstly, more patches of natural, minimally disturbed vegetation need to be sampled to determine the GEDI-derived expected vegetation profile of natural forest across the ecoregion, so that it can be used as a baseline. More sampling is also required along the gradient of land use intensity in order to estimate the shape of the continuous response curves of community composition to habitat degradation. Regarding the use of bioacoustics, it is required to experimentally confirm the assumption that the estimated functional signatures of avian communities are robust to various biases associated with the limitations of bioacoustics technology. It would also be important to investigate whether it might be possible to optimise the use of the ARUs so that they use less energy, in order to limit the decreasing recording effort with increasing vegetation density. For example, it might be possible to robustly estimate functional signatures by only turning on the recorders during a certain time of day (de Araujo et al., 2021) or at a certain temporal frequency (Wood et al., 2020).

Patches of forest, low-intensity, and high-intensity agriculture are not isolated systems as they are embedded into agro-ecological landscapes. In addition to estimating the shape of the continuous habitat-biodiversity relationship, designing biodiversity-friendly productive landscapes in tropical regions requires taking into account the spatial context and configuration (Bennett et al., 2020; Valente et al., 2022). While patches of protected natural forest are crucial for the conservation of specialist species, as highlighted in our results, productive landscapes in the tropics have typically few natural forests remaining, and biodiversity-friendly agricultural practices, such as shade-grown coffee and cocoa, could play a critical conservation role as a refuge for biodiversity in areas that have been hard-hit by deforestation (Perfecto et al., 1996) but also a high-quality matrix connecting patches of natural forest (Perfecto & Vandermeer, 2010; Estrada-Carmona et al., 2022). The monitoring approach that we propose in this study, combining remote sensing with the automated analysis of PAM data to estimate habitat quality and avian community functional signatures as indicators, could be used across a spatial context based on statistical matching techniques (Schleicher et al., 2019) and combined with connectivity theory (Albert et al., 2017), to guide the design of biodiversity-friendly productive tropical landscapes. This way, it could be used to contribute to the ongoing monitoring efforts of the Kunming-Montreal Global Biodiversity Framework to stem the decline of tropical biodiversity.

## Acknowledgements

This work was made possible by support from the Wolfson Foundation and UCL Grand Challenges to M.S.

## Author contributions

M.S. conceived and designed the study. M.S., J.G-H., N.F., S.S., V.C., C.O., F.G.G. and G.L.B. conducted the data collection and data processing. M.S., J.G-H. and N.F. performed the analysis. S.S., M.D., J.V. and I.P. provided critical support and assistance for the collection, analysis, and interpretation of data. M.S. drafted the manuscript. All authors contributed critical input to the manuscript.

## Data accessibility statement

The spaceborne LiDAR data from GEDI is open-access and free to use for research. The raw audio data and handheld LiDAR data is available upon request. Data for sampling effort, species detections obtained from BirdNET, and habitat quality metric obtained from processing LiDAR data, together with the computer code used for the analysis, are available at: https://github.com/msomveille/coffee_birds.git

## Supplementary Material

**Figure S1:**
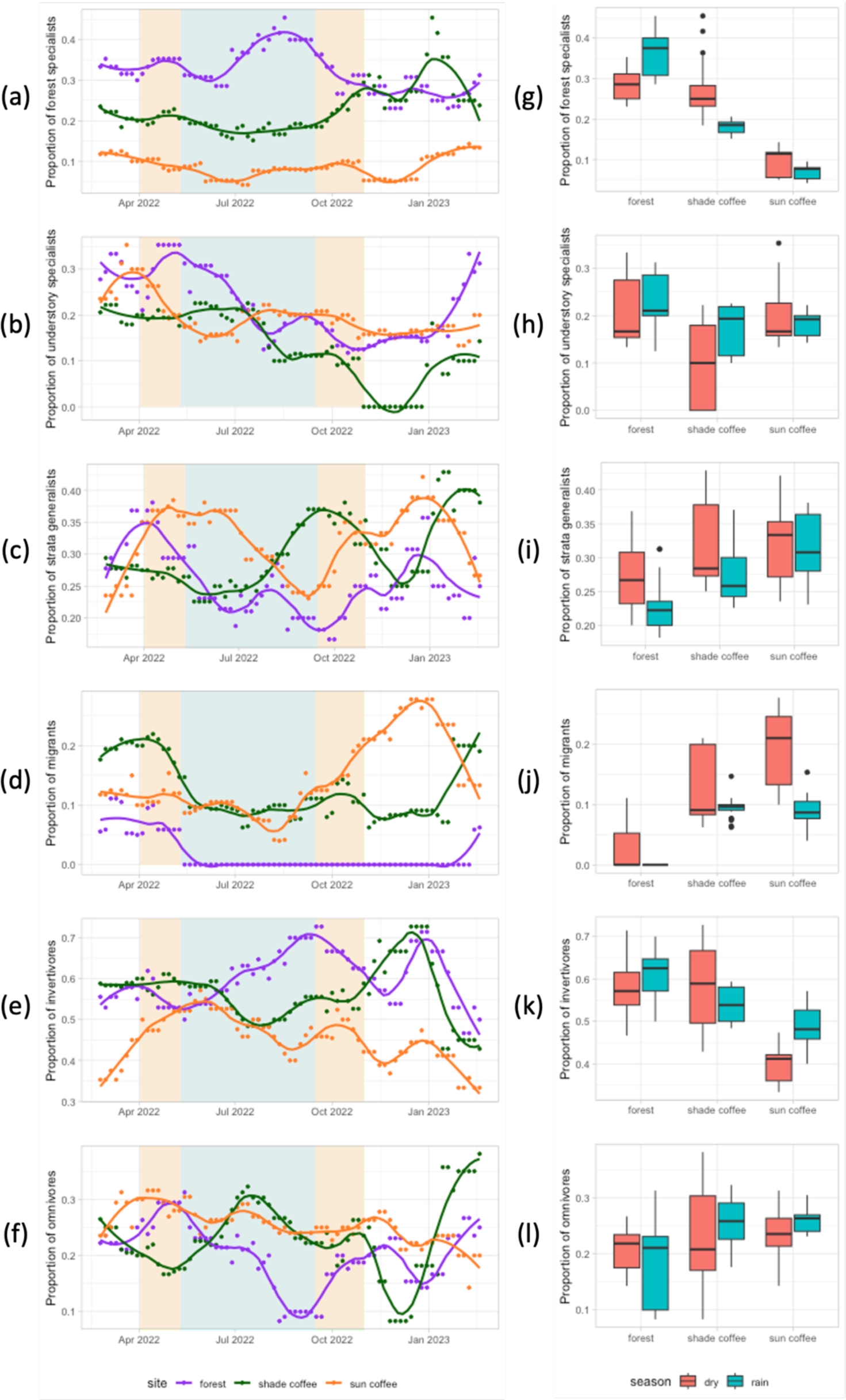
Annual trends in functional signatures for a BirdNET precision threshold of 0.7, as part of the sensitivity analysis. Functional signatures per 5-day period throughout the annual cycle (a–f) and per season (g–l) for each site (shade coffee, sun coffee, and forest), showing trends in the proportion of (a, g) forest specialists, (b, h) understory specialists, (c, i) strata generalists, (d, j) migrants, (e, k) invertivores, and (f, l) omnivores. Lines indicate smoothing curves fitted using local polynomial regression with a smoothing parameter of α = 0.25.

**Figure S2:**
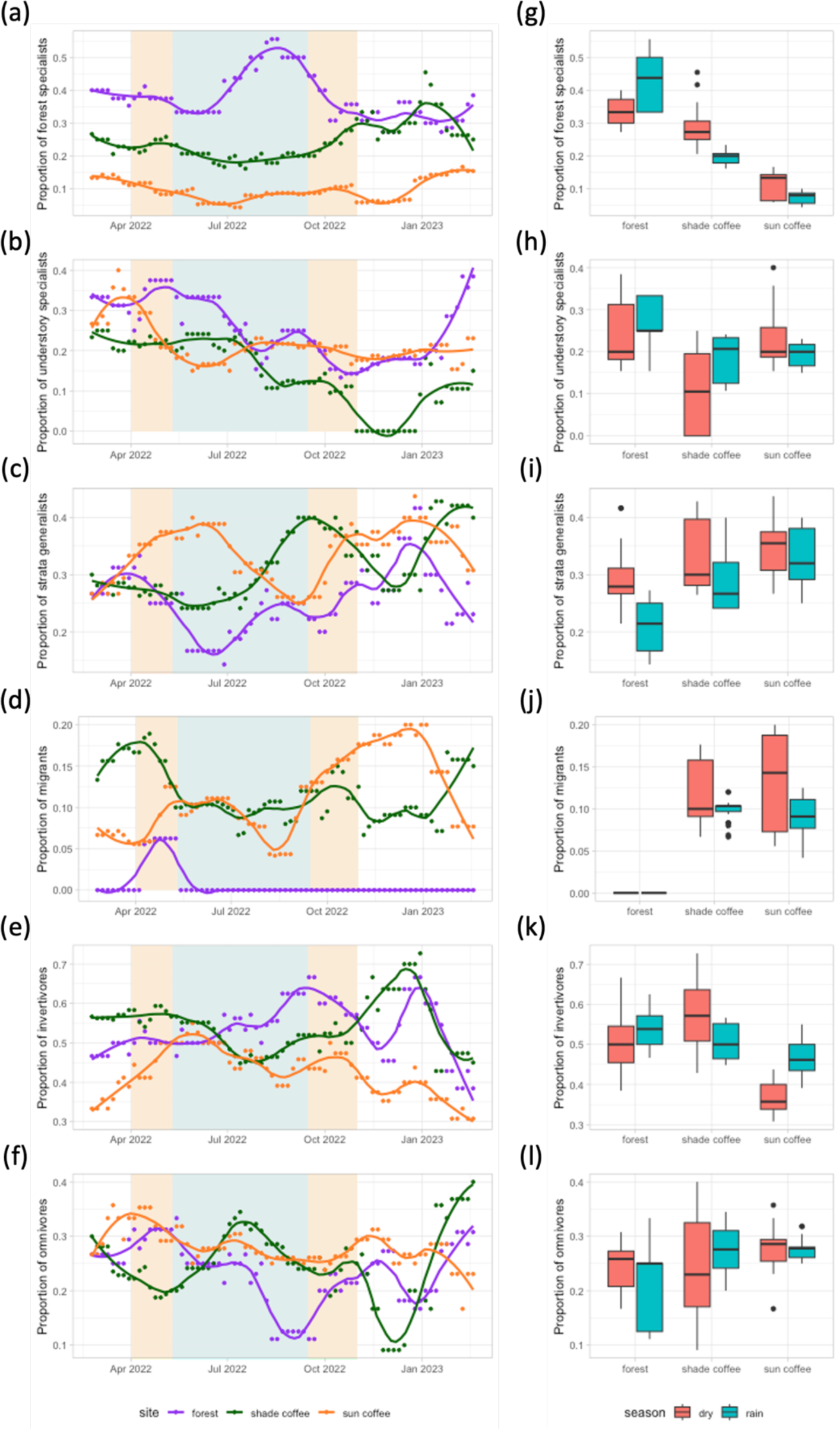
Annual trends in functional signatures for a BirdNET precision threshold of 0.9, as part of the sensitivity analysis. Functional signatures per 5-day period throughout the annual cycle (a–f) and per season (g–l) for each site (shade coffee, sun coffee, and forest), showing trends in the proportion of (a, g) forest specialists, (b, h) understory specialists, (c, i) strata generalists, (d, j) migrants, (e, k) invertivores, and (f, l) omnivores. Lines indicate smoothing curves fitted using local polynomial regression with a smoothing parameter of α = 0.25.

**Figure S3:**
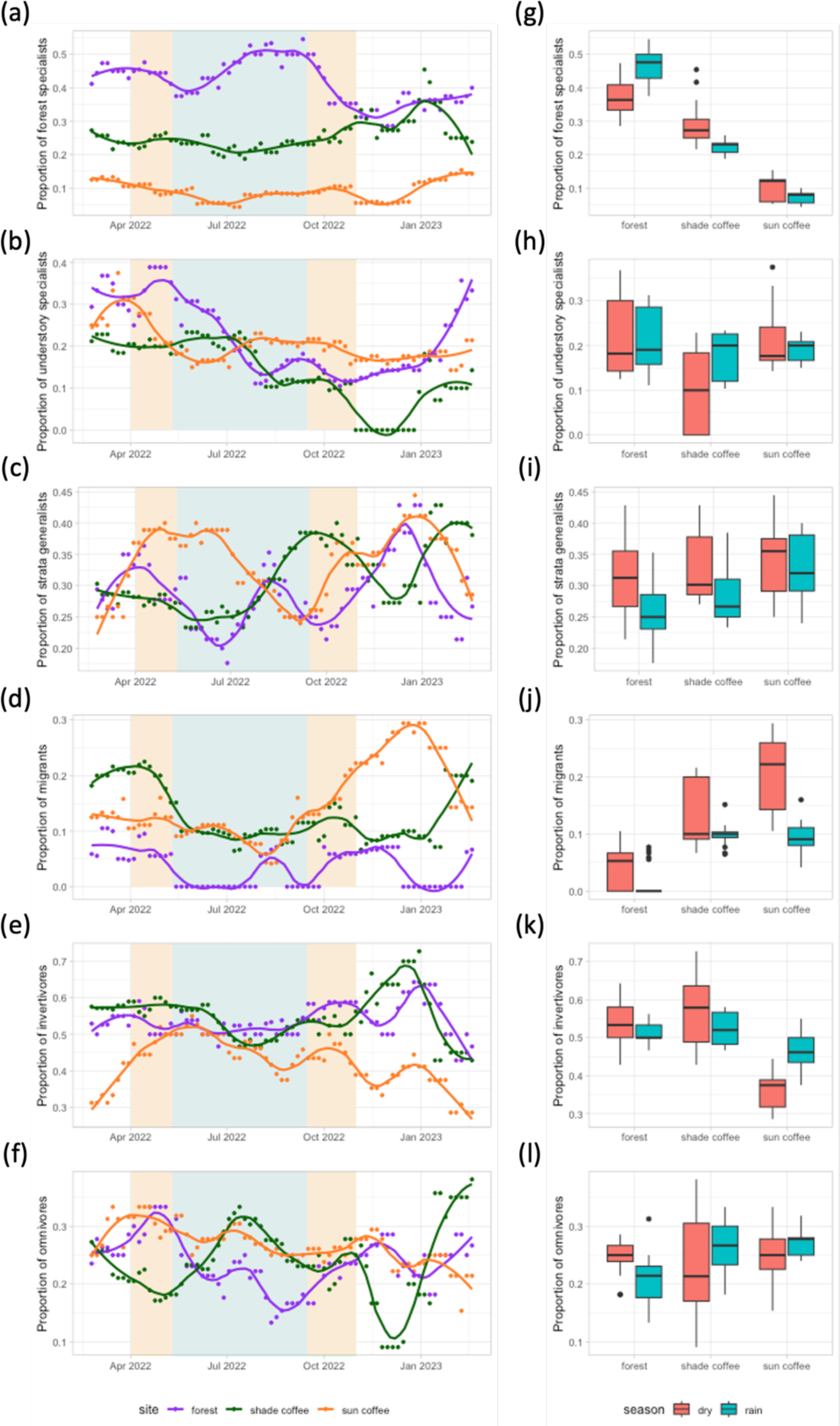
Annual trends in functional signatures for a minimum of 10 detections per species over the whole annual cycle at a site, as part of the sensitivity analysis. Functional signatures per 5-day period throughout the annual cycle (a–f) and per season (g–l) for each site (shade coffee, sun coffee, and forest), showing trends in the proportion of (a, g) forest specialists, (b, h) understory specialists, (c, i) strata generalists, (d, j) migrants, (e, k) invertivores, and (f, l) omnivores. Lines indicate smoothing curves fitted using local polynomial regression with a smoothing parameter of α = 0.25.

**Figure S4:**
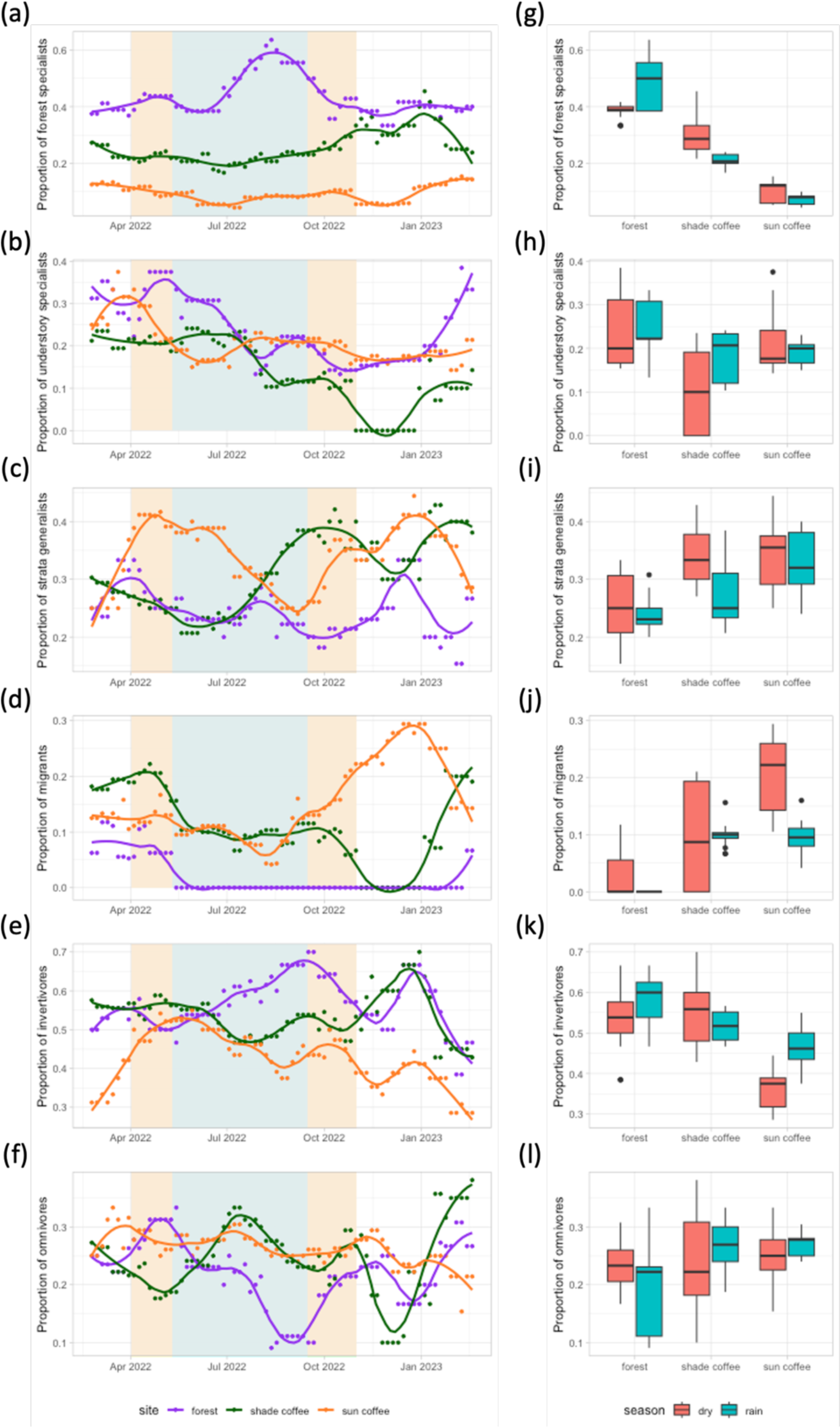
Annual trends in functional signatures for a minimum of 30 detections per species over the whole annual cycle at a site, as part of the sensitivity analysis. Functional signatures per 5-day period throughout the annual cycle (a–f) and per season (g–l) for each site (shade coffee, sun coffee, and forest), showing trends in the proportion of (a, g) forest specialists, (b, h) understory specialists, (c, i) strata generalists, (d, j) migrants, (e, k) invertivores, and (f, l) omnivores. Lines indicate smoothing curves fitted using local polynomial regression with a smoothing parameter of α = 0.25.

**Figure S5:**
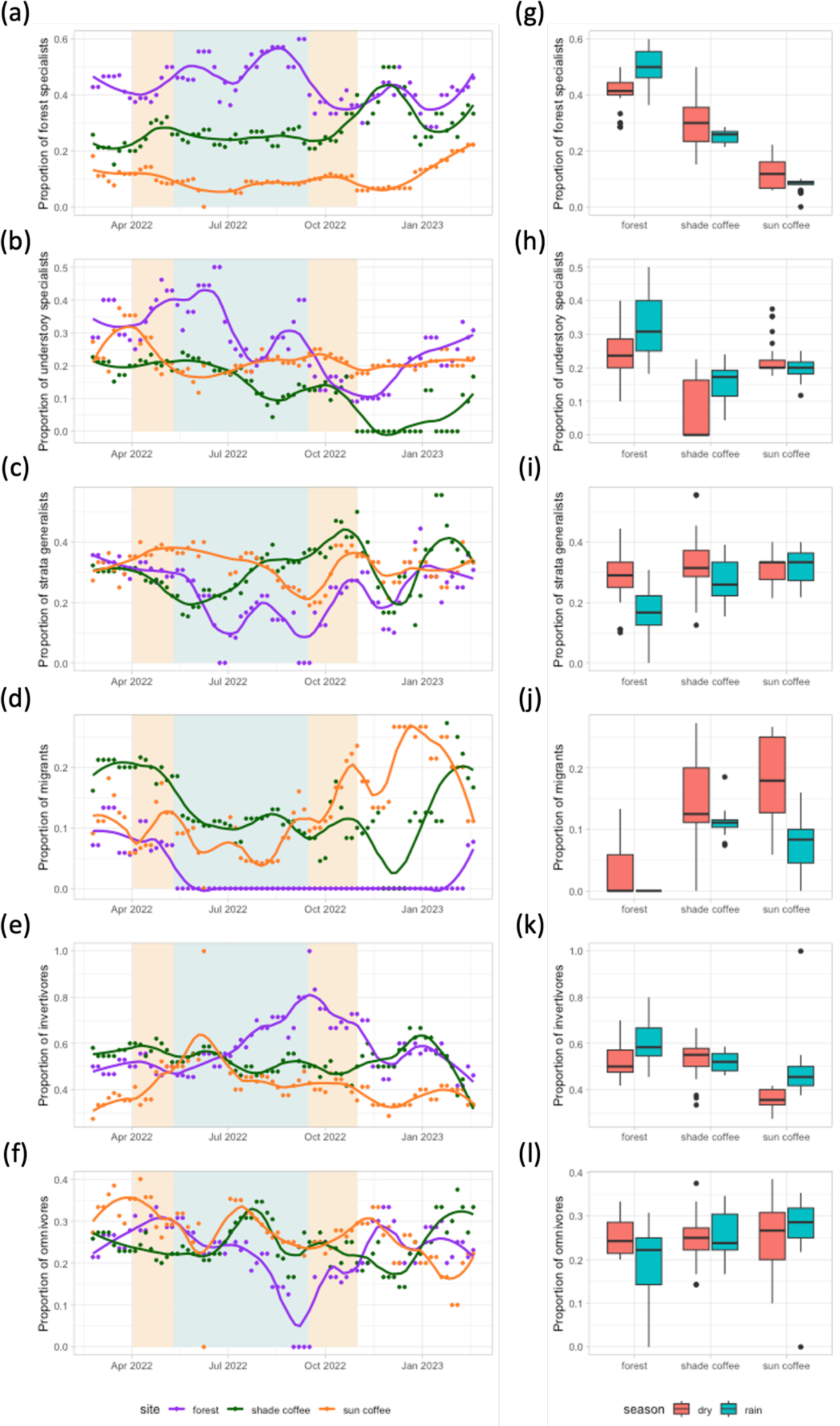
Annual trends in functional signatures for a false absence maximum time span of 20 days, as part of the sensitivity analysis. Functional signatures per 5-day period throughout the annual cycle (a–f) and per season (g–l) for each site (shade coffee, sun coffee, and forest), showing trends in the proportion of (a, g) forest specialists, (b, h) understory specialists, (c, i) strata generalists, (d, j) migrants, (e, k) invertivores, and (f, l) omnivores. Lines indicate smoothing curves fitted using local polynomial regression with a smoothing parameter of α = 0.25.

**Figure S6:**
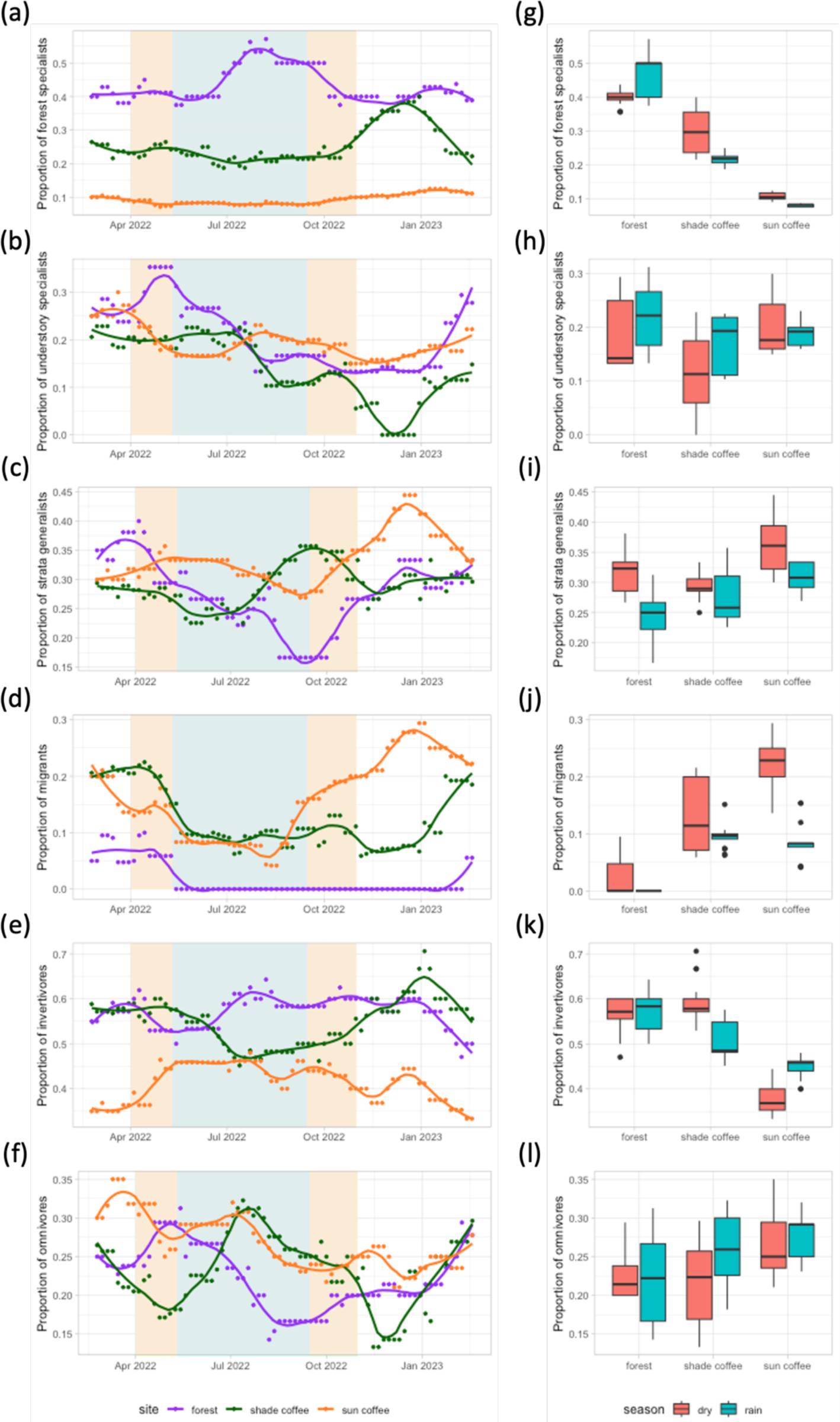
Annual trends in functional signatures for a false absence maximum time span of 60 days, as part of the sensitivity analysis. Functional signatures per 5-day period throughout the annual cycle (a–f) and per season (g–l) for each site (shade coffee, sun coffee, and forest), showing trends in the proportion of (a, g) forest specialists, (b, h) understory specialists, (c, i) strata generalists, (d, j) migrants, (e, k) invertivores, and (f, l) omnivores. Lines indicate smoothing curves fitted using local polynomial regression with a smoothing parameter of α = 0.25.

**Figure S7:**
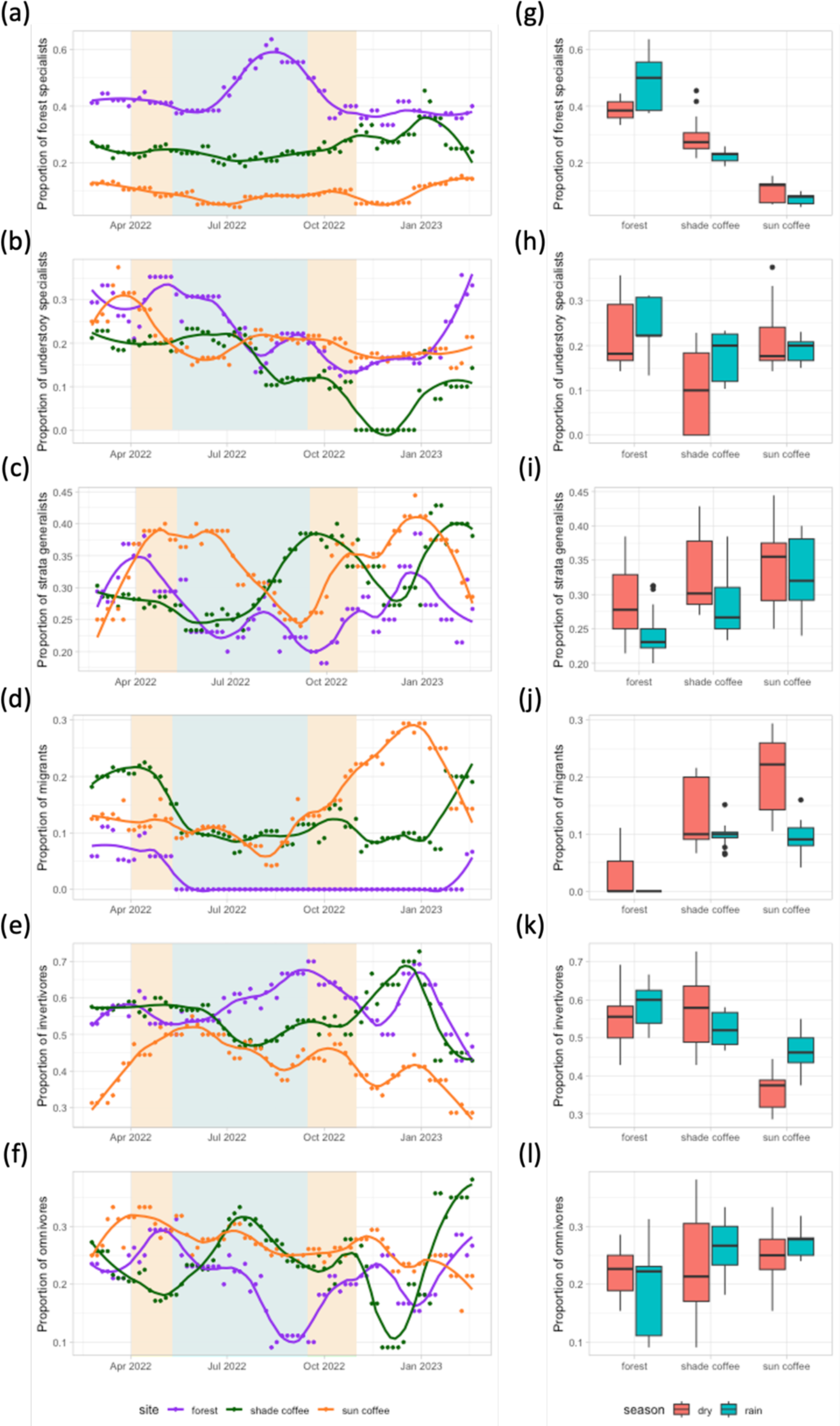
Annual trends in functional signatures for a false presence maximum separation period of 10 days, as part of the sensitivity analysis. Functional signatures per 5-day period throughout the annual cycle (a–f) and per season (g–l) for each site (shade coffee, sun coffee, and forest), showing trends in the proportion of (a, g) forest specialists, (b, h) understory specialists, (c, i) strata generalists, (d, j) migrants, (e, k) invertivores, and (f, l) omnivores. Lines indicate smoothing curves fitted using local polynomial regression with a smoothing parameter of α = 0.25.

**Figure S8:**
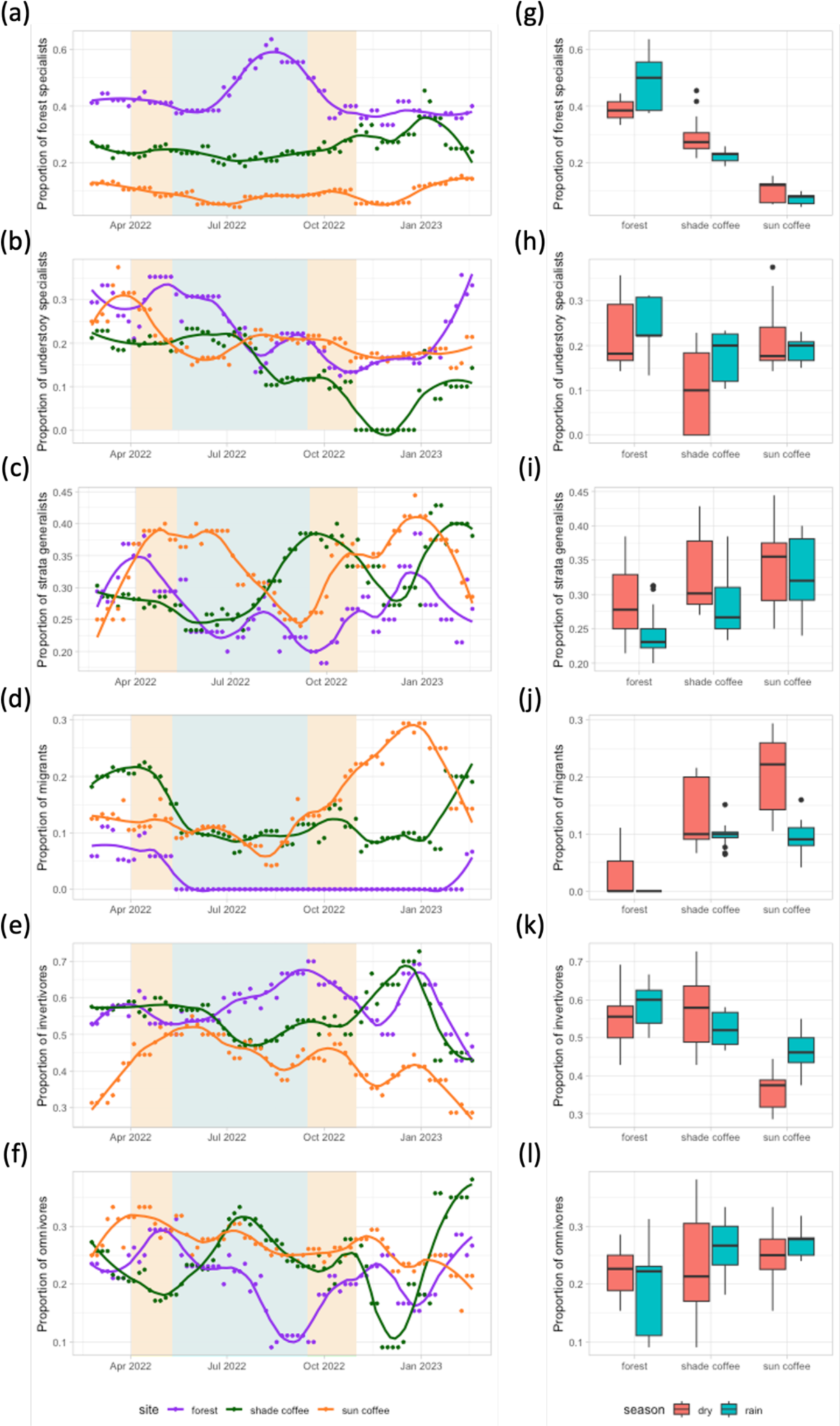
Annual trends in functional signatures for a false presence maximum separation period of 40 days, as part of the sensitivity analysis. Functional signatures per 5-day period throughout the annual cycle (a–f) and per season (g–l) for each site (shade coffee, sun coffee, and forest), showing trends in the proportion of (a, g) forest specialists, (b, h) understory specialists, (c, i) strata generalists, (d, j) migrants, (e, k) invertivores, and (f, l) omnivores. Lines indicate smoothing curves fitted using local polynomial regression with a smoothing parameter of α = 0.25.

## Notes

https://github.com/msomveille/coffee_birds.git

### Competing Interest Statement

The authors have declared no competing interest.

